# The Role of Cilia in the Development, Survival, and Regeneration of Hair Cells

**DOI:** 10.1101/2024.04.01.587636

**Authors:** Hope Boldizar, Amanda Friedman, Tess Stanley, María Padilla, Jennifer Galdieri, Arielle Sclar, Tamara M. Stawicki

## Abstract

Mutations impacting cilia genes lead to a class of human diseases known as ciliopathies. This is due to the role of cilia in the development, survival, and regeneration of many cell types. We investigated the extent to which disrupting cilia impacted these processes in hair cells. We found that mutations in two intraflagellar transport (IFT) genes, *ift88* and *dync2h1,* which lead to the loss of kinocilia, caused increased hair cell apoptosis in the zebrafish lateral line. IFT gene mutants also have a decreased mitochondrial membrane potential, and blocking the mitochondrial uniporter causes a loss of hair cells in wild-type zebrafish but not mutants, suggesting mitochondria dysfunction may contribute to the apoptosis seen in these mutants. These mutants also showed decreased proliferation during hair cell regeneration, but did not show consistent changes in support cell number or proliferation during hair cell development. These results show that disruption of the cilia through either mutations in anterograde or retrograde IFT genes appear to impact hair cell survival but not necessarily development in the lateral line.

## INTRODUCTION

Primary cilia are microtubule-based organelles protruding from cells that serve as localized signaling environments. Mutations in genes impacting cilia lead to a broad class of diseases, known as ciliopathies, that can impact multiple organ systems (Reviewed in Mill et al., 2023; Reiter & Leroux, 2017). One class of genes that has a particularly high impact on cilia is intraflagellar transport (IFT) genes. These are genes that are important for the transport of proteins in cilia and thus also for the formation and maintenance of cilia (Reviewed in Taschner & Lorentzen, 2016). At a cellular level, one of the defects commonly seen in IFT and a subset of other cilia gene mutants is a decrease in cell number. This is seen in several different cell types with the causes for this decreased cell number varying by cell type.

In some cell types, there is evidence of cell death in IFT mutants. Disruption of multiple anterograde IFT genes, through knockdown or mutation, leads to increased apoptosis in photoreceptors (Doerre and Malicki, 2002; Gross et al., 2005; Hudak et al., 2010; Lin-Jones et al., 2003; Lopes et al., 2010; Pazour et al., 2002; Tsujikawa and Malicki, 2004). It is believed that this photoreceptor death is due to the mislocalization of opsin (Doerre and Malicki, 2002; Hudak et al., 2010; Lopes et al., 2010; Pazour et al., 2002; Tsujikawa and Malicki, 2004). Mutation in the retrograde IFT gene, *ift122*, also led to opsin mislocalization, but photoreceptor death was slower than that seen in anterograde IFT mutants (Boubakri et al., 2016). In contrast to this, knockdown of various dynein subunits that serve as retrograde IFT motors did not lead to opsin mislocalization or photoreceptor death (Krock et al., 2009).

There is also evidence of increased apoptosis in the developing nervous system in response to mutations in either IFT or transition zone cilia genes (Abrams and Reiter, 2021; Gorivodsky et al., 2009; Wang et al., 2018). In these cases, it is believed the apoptosis is due to disruptions in either sonic hedgehog (Shh) or Akt signaling.

Nonneuronal cells also show increased cell death when cilia are disrupted, including intervertebral disc cells, thyroid follicular cells, and thyroid cancer cells (Lee et al., 2021; Li et al., 2020a). In thyroid cancer cells this cell death may be due to mitochondria dysfunction. Knockdown of cilia genes in thyroid cancer cells leads to altered mitochondria morphology, decreased mitochondria respiration and ATP production, and decreased mitochondrial membrane potential (Lee et al., 2018; Lee et al., 2021).

Additionally, cilia gene knockdown leads to the oligomerization of VDAC1, which can result in cytochrome c release, and VDAC1 inhibition can block the increased apoptosis normally seen in cilia-depleted thyroid cancer cells (Lee et al., 2021). Decreased mitochondria respiration and ATP production are also seen in kidney proximal tubule cells following *Ift88* knockdown (Fujii et al., 2021), and neurons are more sensitive to mitochondrial inhibitor-induced apoptosis following *Ift88* knockdown (Bae et al., 2019).

In other cell types decreases in cell number seen in IFT mutants are not due to increased cell death, but rather decreased proliferation during development. This is true of neurons in multiple brain regions including the cerebellum, hippocampus, and cortex (Amador-Arjona et al., 2011; Breunig et al., 2008; Chizhikov et al., 2007; Han et al., 2008; Pruski et al., 2019; Spassky et al., 2008; Tong et al., 2014). Decreases in cell proliferation are seen in cilia mutants in other cell types as well including Müller glia cells, oligodendrocyte precursors, chondrocytes, outer annulus fibroblasts cells, osteoblasts, embryonic fibroblasts, and dental pulp stem cells (Cullen et al., 2021; Ferraro et al., 2015; Kitami et al., 2019; Li et al., 2020a; Noda et al., 2016; Pruski et al., 2019; Song et al., 2007; Tao et al., 2023; Yuan et al., 2019).

In addition to their role in controlling cell number developmentally cilia have also been implicated in the regeneration of numerous cell types. Knockdown of *Ift20* decreases proliferation and dedifferentiation of Müller glia cells (Ferraro et al., 2015) and loss of *Ift88* in muscle stem cells leads to decreased proliferation and regenerative ability of muscle (Palla et al., 2022). Regeneration in the heart is also reduced when cilia are disrupted, due to a loss of notch signaling (Li et al., 2020b). Cilia have also been shown to be important for normal notch signaling in epidermal keratinocytes, corneal epithelium cells, and hematopoietic stem and progenitor cells (Ezratty et al., 2011; Grisanti et al., 2016; Liu et al., 2019).

Hair cells contain a single primary cilium known as the kinocilium. It has previously been shown that mutations in genes that lead to the loss of the kinocilium result in a reduction of hair cells (Grati et al., 2015; Stawicki et al., 2016; Stawicki et al., 2019; Tsujikawa and Malicki, 2004). At least part of this decreased hair cell number is likely caused by hair cell death as there is evidence of increased apoptosis in the inner ear of *ift88* zebrafish mutants (Blanco-Sánchez et al., 2014; Tsujikawa and Malicki, 2004), along with evidence of the mistrafficking of proteins in these mutants similar to what is seen in photoreceptors when anterior IFT genes are disrupted (Blanco-Sánchez et al., 2014). However, to our knowledge, no one has ever looked at proliferation during hair cell development or regeneration in these mutants, nor has the reason for reduced hair cell number in retrograde IFT mutants which don’t show protein trafficking defects (Stawicki et al., 2019) been investigated. Therefore, we wished to more thoroughly investigate how mutations in both anterograde and retrograde IFT mutants impacted hair cell number. We carried out this work in the zebrafish lateral line which is a system of surface-localized hair cells allowing for easy access to these cells *in vivo*.

We found evidence of apoptosis in mutants of both the anterior IFT gene, *ift88,* and retrograde IFT gene, *dync2h1* in lateral line hair cells. However, the number of apoptotic cells observed did not fully account for the hair cell number differences seen in these mutants. We also found that mutations in IFT genes led to a slight reduction in mitochondrial membrane potential and that blocking mitochondrial calcium uptake decreased hair cell number in wild-type but not mutant zebrafish. Hair cell development did not seem as impacted as hair cell survival. We did not see alterations in supporting cell number in either mutant. Likewise, proliferation during hair cell development was not consistently disrupted in both mutants, though was we did see a reduction of proliferative cells in *ift88* mutants in some situations. In contrast to this, we observed decreases in proliferation during hair cell regeneration in both mutants and a slight decrease in the number of hair cells that were regenerated. However, it is not clear whether these defects were due to impacts on regeneration itself or the smaller number of initial hair cells in these mutants. Taken together these results suggest that both anterograde and retrograde IFT genes impact hair cell survival and potentially regeneration, but not necessarily hair cell development.

## METHODS

### Animals

All experiments were carried out using fish from either the *dync2h1^w46^* or *ift88^tz288^*mutant lines (Doerre and Malicki, 2002; Stawicki et al., 2016). Mutant alleles were maintained in heterozygous animals in the *AB wild-type background and these heterozygotes were incrossed to get larvae for experiments. Wild-type siblings and mutants were separated based on body morphology phenotypes (Ryan et al., 2013; Tsujikawa and Malicki, 2004). Fish larvae were raised in embryo media (EM) consisting of 1 mM MgSO_4_, 150 μM KH_2_PO_4_, 42 μM Na_2_HPO_4_, 1 mM CaCl_2_, 500 μM KCl, 15 mM NaCl, and 714 μM NaHCO_3._ They were housed in an incubator maintained at 28.5°C with a 14/10 hour light/dark cycle. The Lafayette College Institution Animal Care and Use Committee approved all experiments.

### Immunostaining

Fixation and antibody staining was carried out as previously described (Stawicki et al., 2014). The following primary antibodies were used, mouse anti-otoferlin (Developmental Studies Hybridoma Bank, HCS-1) diluted at 1:100, rabbit anti-cleaved caspase-3 (Cell Signaling Technology, 9661) diluted at 1:250, and rabbit anti-Sox2 (GeneTex, GTX124477). Additionally, for EdU and Caspase experiments nuclei were labeled with DAPI (MilliporeSigma, D9542) at 2μg/ml diluted in PBS. Fish were incubated in this solution for 10 minutes as their last wash out of secondary antibody.

### Cell Counts

Hair cell and cleaved caspase-3 positive cells were counted using an Accu-Scope EXC-350 microscope under the 40X objective. For most experiments the posterior lateral line neuromasts, P1-P9 were counted (Alexandre and Ghysen, 1999). For the RU360 experiments the following anterior lateral line neuromasts were counted OP1, M2, IO4, O2, MI2 and MI1 (Raible and Kruse, 2000). In both cases for hair cell counts the total number of hair cells counted for each fish were divided by the number of neuromasts counted to get an average hair cell/neuromast number for each fish. For the cleaved caspase counts data is shown as the total number of cells counted for each fish due to the small number. Representative images were imaged on a Zeiss LSM800 confocal microscope using the Zen Blue software using a 40x water immersion objective. Z-stacks were taken consisting of 15 (caspase) or 12 (regeneration hair cells) slices separated by 1μm and then maximum projection images were made in Fiji.

### JC-1 Labeling and Analysis

JC-1 was used as previously described (Pickett et al., 2018). Fish were incubated with 1.5 μM JC-1 (ThermoFisher, T3168) diluted in EM for 30 minutes, washed 3x with plain EM and then left in the third EM wash for 90 minutes. After 90 minutes fish were anesthetized with MS-222 and imaged on a Zeiss LSM800 using a 40x water immersion objective. For imaging both the green and red signals were obtained using a 488 nm laser. The green signal looked at 410-546 nm emissions whereas the red signal looked at 585-700 nm emissions. One neuromast was imaged for each fish. A z-stack was taken consisting of 5 slices separated by 1 μm. Image analysis was carried out in Fiji. Maximum projections were made and then the average red and green fluorescence was measured. From these numbers a red/green ratio was generated. For each mutant data from two separate experiments were combined. Thus, for each individual experiment the data was normalized to the average red/green ratio of the wild-type siblings for that experiment.

### Drug Treatment

For all drug treatments fish were loaded into netwell inserts in 6 well plates containing EM. The netwell inserts containing the fish were then moved into new 6 well plates containing the drugs being used for treatment diluted in EM. Drugs used were either 500 nM RU360 (MilliporeSigma, 57440) or 200 μM Gentamicin (MilliporeSigma, G1272). Following drug treatment fish were washed three times with plain EM, again by moving netwell inserts to new 6 well plates containing plain EM.

### EdU Labeling and Analysis

EdU labeling was carried out similar to what has previously been described (Thomas and Raible, 2019). Fish were incubated in EdU (ThermoFisher, A10044 or C10340) at 500 μM for 24 hours. After fixation EdU positive cells were labeled via Click-iT reaction using the Click-iT^TM^ EdU cell proliferation kit for imaging (ThermoFisher, C10340) following the protocol that came with the kit. Hair cells and nuclei were then labeled as mentioned above in immunostaining. To count the number of EdU positive hair cells, fish were imaged on a Zeiss LSM800 microscope. A z-stack was taken consisting of 15 (developmental) or 12 (regeneration) slices separated by 1μm. Image analysis was carried out in Fiji. Composite images were generated combining the EdU, Otoferlin and DAPI labels and then the number of EdU positive hair cells were manually counted in Fiji using the cell counter tool. For each fish the P1, P2, and P3 neuromasts were counted and then an average EdU positive hair cell/neuromast number was generated. Counts were carried out using the full z-stacks, however, the representative images shown are maximum projections which were also made in Fiji.

### Support Cell Number Quantification

Fish were fixed at 6 dpf and hair cells and support cells were labeled using the Otoferlin and Sox2 antibody respectively as mentioned above in immunostaining. To count the number of support cells, fish were imaged on a Zeiss LSM800 microscope. A z-stack was taken consisting of slices separated by 1μm through the entirety of each neuromast. Image analysis was carried out in Fiji. Composite images were generated combining the Sox2 and otoferlin labels and then the number of Sox2 positive cells were manually counted in Fiji using the cell counter tool. For each fish the P1, P2, and P3 neuromasts were counted and then an average Sox2 positive support cell/neuromast number was generated. Counts were carried out using the full z-stacks, however, the representative images shown are maximum projections which were also made in Fiji.

### Statistics and Graphing of Data

Most statistics were calculated in GraphPad Prism 6 with the exception of the 3-way ANOVA for the hair cell regeneration data which was calculated in SPSS version 29. The specific statistical tests used for each experiment are mentioned in the figure legends. The graphs were all generated in GraphPad Prism 6. Graphs show either individual data points along with the averages and standard deviations as superimposed lines, with the exception of the regeneration hair cell counts which are shown as bar graphs showing the average and standard deviations for simplicity’s sake given the number of groups.

## RESULTS

### Lateral line hair cells of IFT gene mutants undergo apoptosis

It has previously been shown that hair cells undergo apoptosis in the inner ear of *ift88* mutants (Blanco-Sánchez et al., 2014; Tsujikawa and Malicki, 2004). We wished to investigate whether apoptosis also occurred in both anterograde and retrograde IFT gene mutants in the lateral line. To do this we stained wild-type siblings and mutants of both the anterograde IFT gene, *ift88,* and the retrograde IFT gene, *dync2h1,* fixed at 4, 5, and 6 days post fertilization (dpf) with an antibody for both the hair cell-specific protein otoferlin and cleaved caspase-3. Similar to what has previously been found (Stawicki et al., 2016) we saw a significant decrease in hair cell number in both IFT gene mutants, particularly at later ages (Figures 1A, B & 2A, B). Additionally, we saw a small number of cleaved caspase-3 positive hair cells in both IFT gene mutants whereas we never saw any in wild-type sibling controls, suggesting that hair cells in these mutants do undergo apoptosis (Figures 1C & 2C). However, the number of cleaved caspase-3 positive cells we observed in the mutants was considerably lower than the difference in hair cell number between wild-type siblings and mutants.

**Figure 1:**
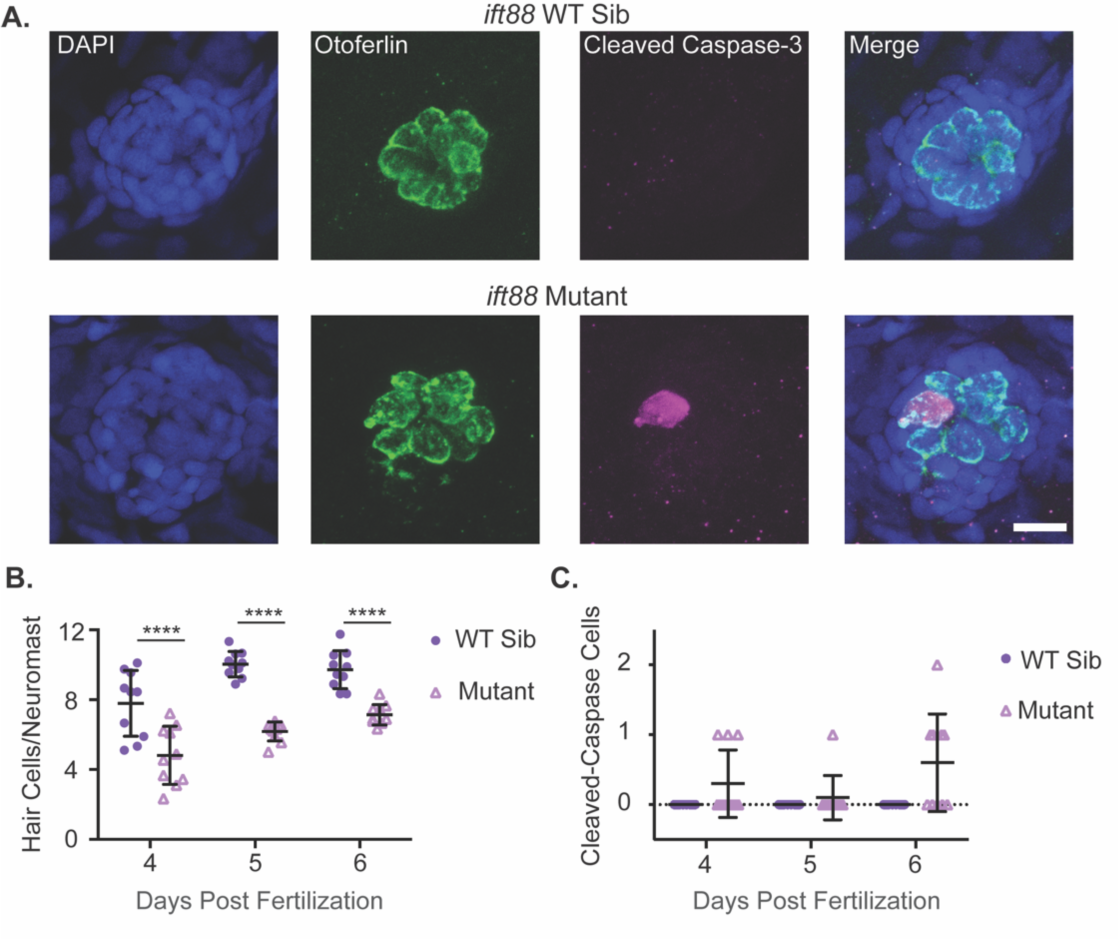
*ift88* mutants show decreased hair cell number and hair cells undergoing apoptosis. (A) Representative images of neuromasts from *ift88* wild-type siblings (top) and *ift88* mutants (bottom) at 6 dpf. Nuclei are labeled in blue with DAPI, hair cells in green with the Otoferlin antibody and cells undergoing apoptosis in magenta with the cleaved caspase-3 antibody. Scale bar = 10μm. (B) Quantification of hair cells/neuromasts in *ift88* wild-type siblings and mutants at 4, 5 and 6 dpf. Genotype and age were significant sources of variation by 2-way ANOVA (p<0.0001), but the interaction between the two variables was not (p = 0.2377). **** = p<0.001 by Šídák’s multiple comparisons test comparing the wild-type siblings and mutants at the different ages. n=10 for each group. (C) Quantification of the total number of cleaved caspase-3 positive hair cells in the 9 neuromasts of the posterior lateral line in *ift88* wild-type siblings and mutants at 4, 5 and 6 dpf. Genotype was a significant source of variation by 2-way ANOVA (p = 0.001) whereas age (p = 0.1089) and the interaction between the two variables (p = 0.0189) were not. n=10 for each group.

**Figure 2:**
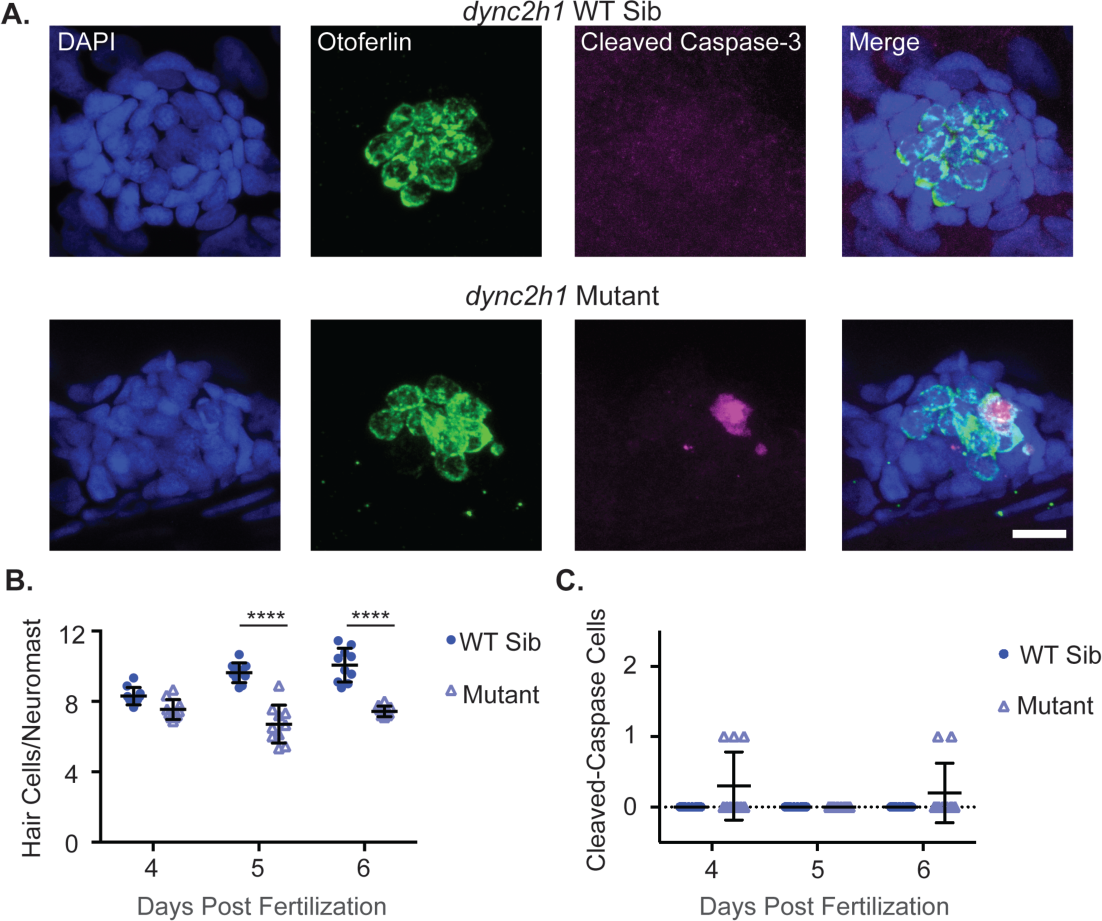
*dync2h1* mutants show decreased hair cell number and hair cells undergoing apoptosis. (A) Representative images of neuromasts from *dync2h1* wild-type siblings (top) and *dync2h1* mutants (bottom) at 6 dpf. Nuclei are labeled in blue with DAPI, hair cells in green with the Otoferlin antibody and cells undergoing apoptosis in magenta with the cleaved caspase-3 antibody. Scale bar = 10μm. (B) Quantification of hair cells/neuromasts in *dync2h1* wild-type siblings and mutants at 4, 5 and 6 dpf. Genotype (p<0.0001), age (p = 0.0003), and the interaction between the two variables (p<0.0001) were significant sources of variation by 2-way ANOVA. **** = p<0.001 by Šídák’s multiple comparisons test comparing the wild-type siblings and mutants at the different ages. n=10 for each group. (C) Quantification of the total number of cleaved caspase-3 positive hair cells in the 9 neuromasts of the posterior lateral line in *dync2h1* wild-type siblings and mutants at 4, 5 and 6 dpf. Genotype was a significant source of variation by 2-way ANOVA (p = 0.0169) whereas age (p = 0.1918) and the interaction between the two variables (p = 0.1918) were not. n=10 for each group.

### Mitochondria dysfunction may play a role in hair cell apoptosis in IFT gene mutants

Mutation or knockdown of cilia genes has been shown to alter mitochondria function in multiple cell types (Fujii et al., 2021; Lee et al., 2018) and in some cases, this may be linked to increased cell death (Bae et al., 2019; Lee et al., 2021). Additionally, mutations of IFT genes in hair cells lead to a reduction in rapid FM1-43 uptake suggesting a decrease in hair cell activity (Stawicki et al., 2016), and it has been shown that reduction in hair cell activity can lead to reduced mitochondrial membrane potential (Pickett et al., 2018). To test if mitochondrial membrane potential was reduced in hair cells of IFT gene mutants we used the ratiometric mitochondrial membrane potential indicator dye, JC-1. JC-1 aggregates in response to mitochondrial membrane polarization, switching from its monomeric green fluorescent form to a red fluorescing form (Reers et al., 1991). Thus, the red/green ratio is influenced by mitochondrial membrane potential with a smaller ratio being indicative of a less polarized mitochondrial membrane potential and vice versa. We found a small but significant decrease in the red/green JC-1 ratio of both IFT gene mutants (Figure 3).

**Figure 3:**
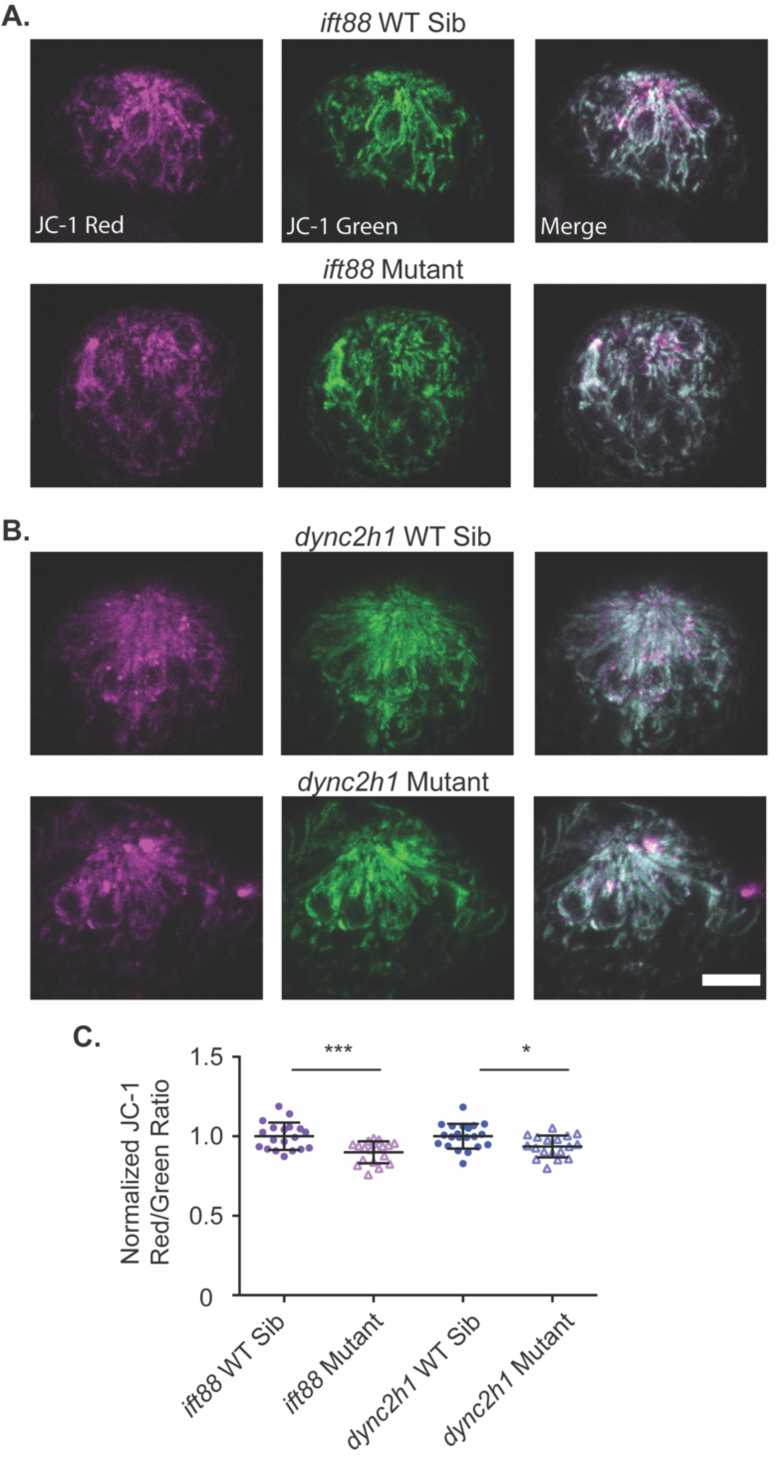
Mitochondrial membrane potential is slightly reduced in IFT gene mutants. (A) Representative images of neuromasts from (A) *ift88* or (B) *dync2h1* wild-type siblings (top) and mutants (bottom) labeled with JC-1. Scale bar = 10μm. (C) Quantification of the JC-1 Red/Green ratio for both *ift88* and *dync2h1* mutants and wild-type siblings. There was a significant difference in both mutants by unpaired Student’s t-test, p = 0.0004 for *ift88,* shown as ***, and p = 0.0105 for *dync2h1,* shown as *. n = 19 wild-type siblings and 18 mutants for *ift88* and n = 21 wild-type siblings and 18 mutants for *dync2h1* mutants. n numbers are not equal for all groups as some images were not usable due to the movement of fish during imaging.

In other hair cell mutants with reduced hair cell number and mitochondrial membrane potential decreases it has been shown that treatment with the mitochondrial calcium uniporter inhibitor RU360 can partially rescue decreases in hair cell number (Santra and Amack, 2021). We wished to test if this was true for IFT gene mutants. To do this fish were treated with RU360 at 500nM from 4-6 dpf and then fish were fixed and hair cells were labeled for counting. Similar to what had previously been found with extended RU360 treatment (Santra and Amack, 2021) we found a significant decrease in the number of hair cells in wild-type siblings. However, we saw no changes in hair cell number in either IFT gene mutant following RU360 treatment (Figure 4). Additionally, we noted that hair cell number in wild-type siblings treated with RU360 was similar to hair cell number in IFT gene mutants in both conditions.

**Figure 4:**
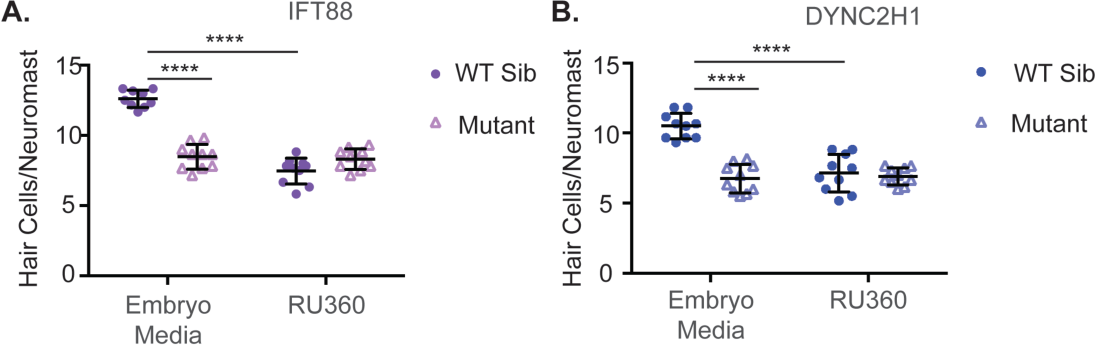
Inhibition of the mitochondrial uniporter decreases hair cell number in wild-type siblings, but not IFT gene mutants. Quantification of hair cells/neuromast in (A) *ift88* and (B) *dync2h1* wild-type siblings and mutants with or without RU360 treatment. In the case of both mutants genotype, drug treatment, and the interaction between the two variables was a significant source of variation by 2-way ANOVA (p<0.0001). **** = p<0.0001 by Šídák’s multiple comparisons test comparing the two groups at the edges of the lines under the asterisks. n = 10 for all groups with the exception of the *ift88* WT sibling EM group where n = 9 due to the loss of fish during the staining process.

### Proliferation during hair cell development is not consistently reduced in IFT gene mutants

We next wanted to test if proliferation was altered during development in cilia gene mutants. To do this we treated fish with EdU from 3-4 dpf to label proliferating cells. We then fixed fish at 5 dpf, labeled hair cells, and counted the number of EdU-positive hair cells in both wild-type siblings and mutants of *ift88* and *dync2h1* mutants. While the average numbers of EdU-positive hair cells were slightly reduced for each mutant, particularly *ift88* mutants, these differences were not statistically significant (Figure 5A-C). As we know hair cells are dying in IFT gene mutants (Figures 1 & 2) we next wanted to test how delaying the fixation point would impact the number of EdU-positive hair cells seen. Therefore, we again treated fish with EdU from 3-4 dpf to label proliferating cells during this period, but then fixed and stained fish at 6dpf to label hair cells. Using this paradigm we found that there now were significantly fewer EdU-positive hair cells in *ift88* gene mutants whereas there continued to be no significant differences in the number of Edu-positive hair cells between the *dync2h1* wild-type siblings and mutants (Figure 5D).

**Figure 5:**
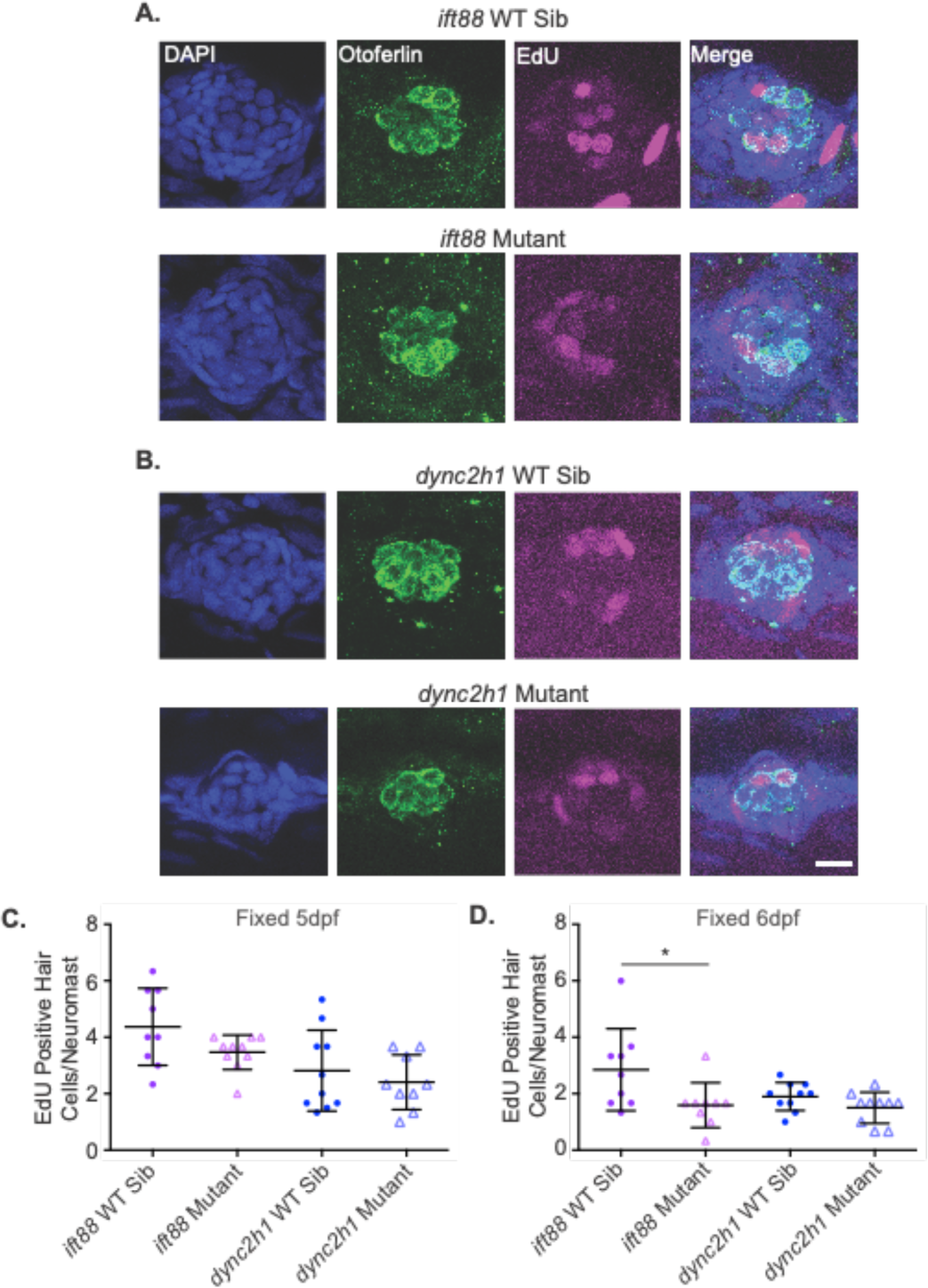
Proliferation during hair cell development is not significantly decreased in both IFT gene mutants. Representative images of neuromasts from (A) *ift88* and (B) *dync2h1* wild-type siblings (top) and mutants (bottom) at 5 dpf. Cells that proliferated from 3-4 dpf are EdU positive as shown in magenta. Nuclei are labeled in blue with DAPI and hair cells in green with the otoferlin antibody. Scale bar = 10μm. (C) Quantification of the number of EdU positive hair cells/neuromast for each mutant when fixed at 5 dpf. There were no significant differences when comparing wild-type siblings to mutants by unpaired t-test for either mutant (p = 0.096 for *ift88* and p = 0.481 for *dync2h1*). For *ift88* Welch’s correction was used due to the unequal variances between wild-type siblings and mutants. n = 10 for *ift88* mutants and *dync2h1* WT siblings and n = 9 for *ift88* WT siblings and *dync2h1* mutants. (D) Quantification of the number of EdU positive hair cells/neuromast for each mutant when fixed at 6 dpf. *ift88* mutants showed a significant reduction in the number of Edu positive hair cells as compared to their wild-type siblings (p = 0.0411) by unpaired t-test. There were still no significant differences when comparing *dync2h1* wild-type siblings to mutants (p = 0.1055). n = 9 for both *ift88* groups and n = 10 for both *dync2h1* groups. n numbers were not equal in all groups due to the loss of fish in the staining process or issues with images.

### Supporting cell number is not altered in IFT gene mutants

In addition to playing a role in proliferation during development cilia can also be important for the differentiation of some cell types (Liu et al., 2019; Song et al., 2007). In the zebrafish lateral line a subset of cells in neuromasts are specified as hair cells with the remaining cells becoming support cells (reviewed in Thomas et al., 2015).

Therefore, defects in hair cell specification may result in alterations in supporting cell number. To test for changes in support cell number we stained 6 dpf zebrafish with a Sox2 antibody to label supporting cells (Hernández et al., 2007). Doing this we failed to see any significant differences between the number of Sox2 positive cells in wild-type versus mutant zebrafish in either *ift88* or *dync2h1* mutants (Figure 6).

**Figure 6:**
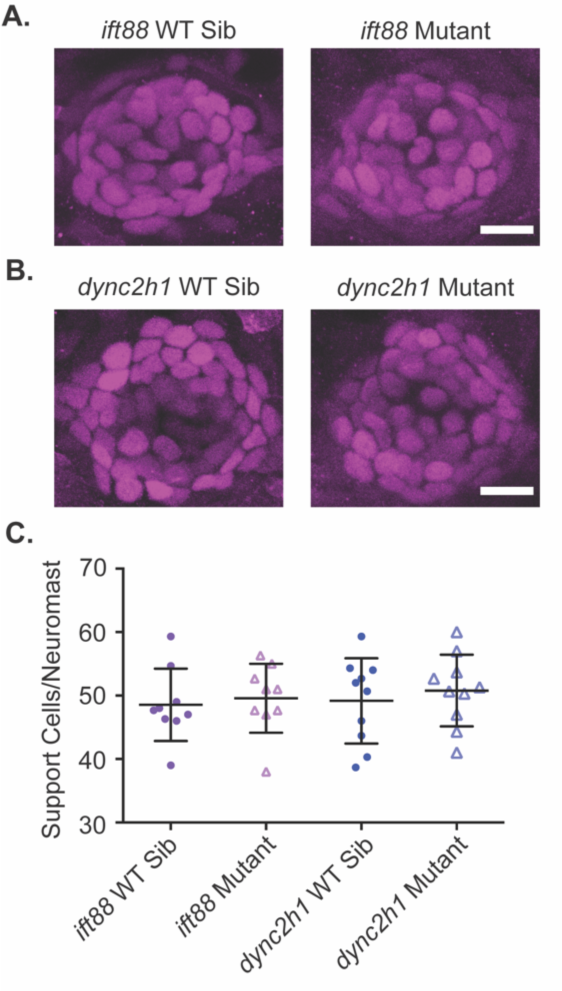
Support cell number is not altered in IFT gene mutants. Representative images of neuromasts from (A) *ift88* and (B) *dync2h1* wild-type siblings (left) and mutants (right) at 6 dpf stained with a Sox2 antibody to label support cells. Scale bar = 10μm. (C) Quantification of the number of Sox2 positive support cells/neuromast for each mutant. There were no significant differences when comparing wild-type siblings to mutants by unpaired t-test for either mutant (p = 0.6982 for *ift88* and p = 0.5632 for *dync2h1*). n = 9 for both *ift88* groups and n = 10 for both *dync2h1* groups. n numbers were not equal in all groups due to issues with images.

### Hair cell regeneration is slightly reduced in IFT gene mutants

Regenerative ability has also been shown to be reduced in some cell types following the loss of cilia (Palla et al., 2022) in some cases due to disruption in notch signaling (Li et al., 2020b). However, in hair cells disruption of notch signaling usually leads to increased regeneration (Daudet et al., 2009; Ma et al., 2008; Mizutari et al., 2013; Warchol et al., 2017). Therefore, we wished to investigate what hair cell regeneration would look like in cilia gene mutants. As IFT gene mutants are strongly resistant to short-term neomycin-induced hair cell death (Stawicki et al., 2016), we used a long-term gentamicin treatment, which they are less resistant to (Stawicki et al., 2019), to trigger hair cell regeneration. Both wild-type siblings and IFT mutant fish were treated with 200 μM gentamicin for 24 hours at 4 dpf after which ½ the fish were fixed to confirm hair cell death and the other ½ were left in plain EM for 48 hours to allow for hair cell regeneration. Another group of fish was kept in plain EM for the first 24 hours and again ½ were fixed and ½ were moved to fresh EM for an additional 48 hours. Both wild-type siblings and IFT mutants experienced significant hair cell death in response to gentamicin treatment (Figure 7). There was also significant hair cell regeneration in both wild-type siblings and IFT mutants in the 48-hour period after gentamicin treatment.

**Figure 7:**
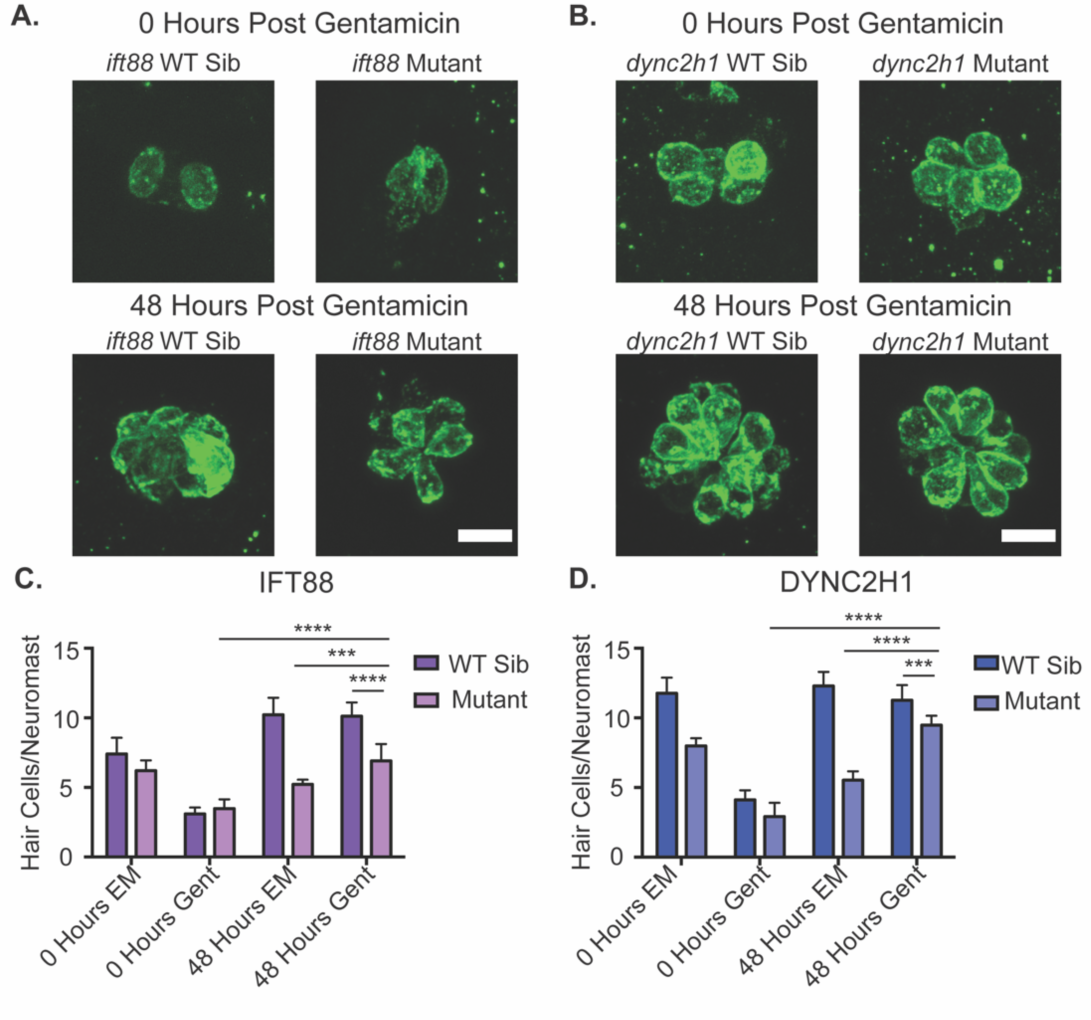
Hair cell regeneration is slightly reduced in IFT gene mutants. Representative images of neuromasts with hair cells labeled using the otoferlin antibody in (A) *ift88* and (B) *dync2h1* wild-type siblings (left) and mutants (right), pre (top) and post (bottom) regeneration. Scale bar = 10μm. Quantification of hair cells/neuromast for (C) *ift88* and (D) *dync2h1* wild-type siblings and mutants during the regeneration experiment. EM groups were fish that were kept in EM the first 24 hours of the experiment whereas Gent groups were fish that were put in 200 μM gentamicin for the first 24 hours of the experiment. 0 hour groups were fixed immediately following those first 24 hours, whereas 48 hour groups were fixed following a 48 hour recovery period in EM. A 3-way ANOVA was carried out for both experiments which found all variables (time, drug, genotype) to be significant sources of variation as well as all two-way interactions between the 3 variables (p<0.001). The three way interaction between variables was found to be significant for the *dync2h1* experiment (p = 0.003), but not the *ift88* experiment (p = 0.796). **** = p<0.0001 and *** = p<0.001 by Šídák’s multiple comparisons test comparing the two groups at the edges of the lines under the asterisks. n = 10 for all groups.

However, in the case of both *ift88* and *dync2h1* mutants, the amount of regeneration was significantly reduced compared to wild-type siblings (Figure 7). Interestingly, despite this reduction in regeneration, the difference in hair cell number between IFT gene mutants and their wild-type siblings was greater in the groups that did not undergo regeneration than those that did. Relatedly, while wild-type siblings showed comparable, if not slightly reduced, hair cell numbers in their 48-hour fix groups that had undergone regeneration groups as compared to the 48-hour fix group that had not undergone regeneration, in both IFT gene mutants there were significantly more hair cells in the 48-hour fix group that had undergone regeneration as compared to the one that had not (Figure 7).

Given the reduction in hair cell regeneration in IFT gene mutants we next wanted to see if proliferation during hair cell regeneration was disrupted in these mutants. To do this fish were again treated with 200 μM gentamicin for 24 hours at 4 dpf, however, this time EdU was included during the gentamicin treatment. Co-treatment of EdU with gentamicin was performed because we found that treating with EdU after gentamicin did not result in the expected increase in EdU-positive hair cells in wild-type siblings treated with gentamicin as compared to the EM group (data not shown). Fish were then washed out of gentamicin and EdU, and given 48 hours to recover in plain EM before they were fixed and labelled with hair cell markers. Both *ift88* and *dync2h1* mutants saw a significant decrease in the number of EdU-positive hair cells as compared to wild-type siblings in the groups that received gentamicin treatment to trigger hair cell regeneration (Figure 8). We also saw a significant decrease in the number of EdU-positive hair cells in the *ift88* mutant EM control group as compared to the *ift88* wild-type sibling EM controls (Figure 8) which is similar to what we previously saw in *ift88* mutants when fish were fixed 48 hours after EdU treatment (Figure 5C).

**Figure 8:**
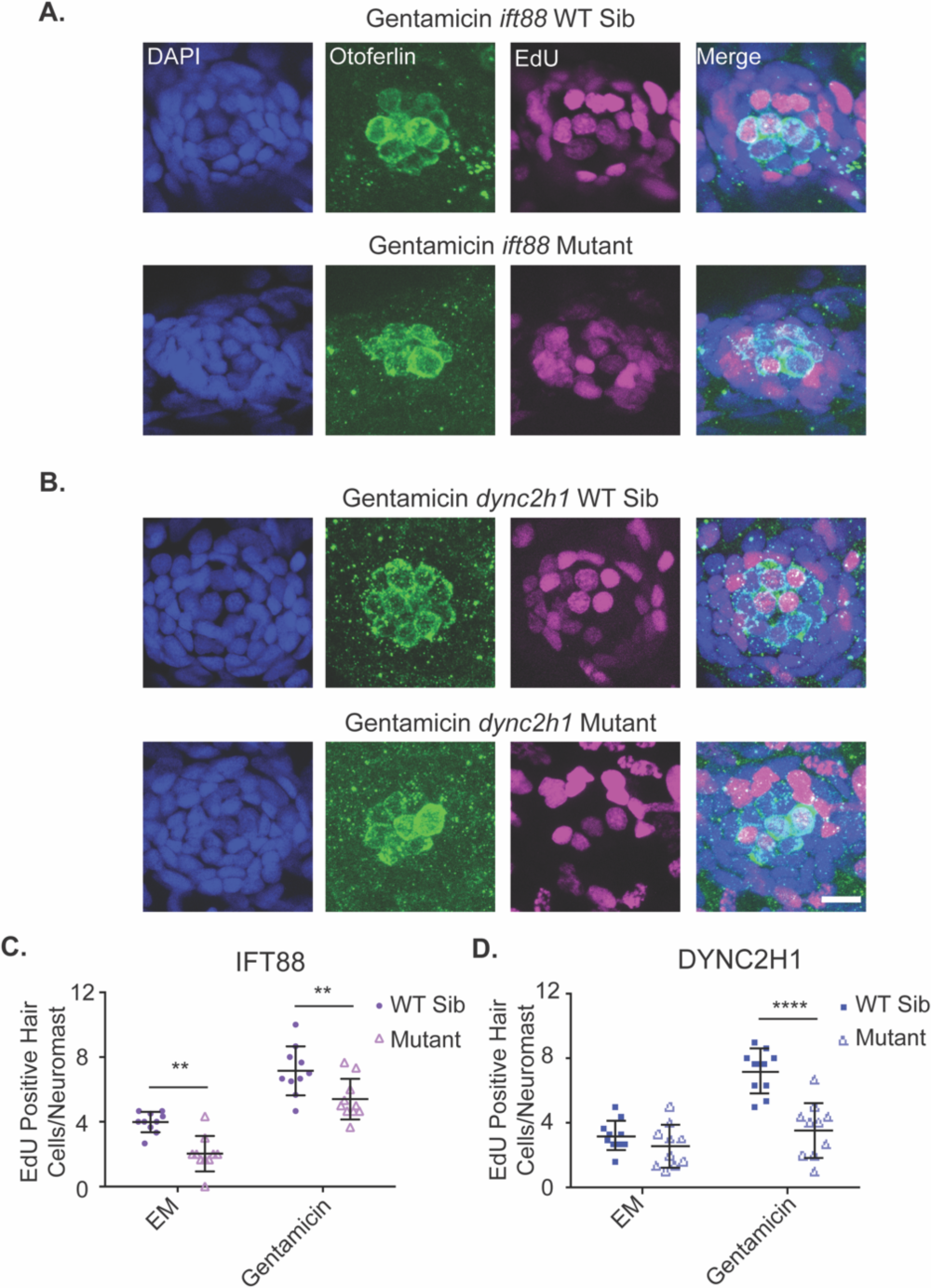
Proliferation during hair cell regeneration is reduced in IFT gene mutants. Representative images of neuromasts from (A) *ift88* and (B) *dync2h1* wild-type siblings (top) and mutants (bottom) at 7 dpf following gentamicin and EdU treatment from 4-5 dpf. Cells that proliferated during the 4-5 dpf time period are EdU positive as shown in magenta. Nuclei are labeled in blue with DAPI and hair cells in green with the otoferlin antibody. Scale bar = 10μm. The graphs show the quantification of EdU positive hair cells/neuromast for (C) *ift88* and (D) *dync2h1* wild-type siblings and mutants 48 hours after either EM or gentamicin treatment. A 2-way ANOVA was carried out for both experiments. For *ift88* both drug treatment and genotype were significant sources of variation (p<0.0001) whereas the interaction between the two variables was not (p=0.7884). For *dync2h1* both of the individual variables were also significant sources of variation (p<0.0001) as was the interaction between the two variables (p=0.0016). ** = p<0.01 and **** = p<0.0001 by Tukey’s multiple comparisons test comparing the two groups at the edges of the lines under the asterisks. n = 10 for all groups.

## DISCUSSION

It has previously been shown that mutations that lead to the loss of kinocilia in hair cells lead to a reduction in hair cell number in both the inner ear and lateral line of zebrafish (Grati et al., 2015; Stawicki et al., 2016; Stawicki et al., 2019; Tsujikawa and Malicki, 2004). In this work, we wished to further investigate the mechanism behind this hair cell number reduction in addition to seeing if there were similar hair cell number changes during hair cell regeneration. Past work has suggested that decreases in hair cell number in the inner ear in mutants of the anterograde IFT gene, *ift88,* are caused by hair cell apoptosis potentially due to the mistrafficking of proteins and subsequent ER stress (Blanco-Sánchez et al., 2014; Tsujikawa and Malicki, 2004). However, it is not obvious that proteins are mistrafficked in mutants of retrograde IFT genes like *dync2h1* (Stawicki et al., 2019). This is similar to what has previously been shown in photoreceptors where anterograde IFT gene mutants show mistrafficking of opsins and subsequent cell death (Doerre and Malicki, 2002; Hudak et al., 2010; Lopes et al., 2010; Pazour et al., 2002; Tsujikawa and Malicki, 2004), however, in retrograde IFT gene mutants there is either no evidence of opsin mistrafficking or cell death (Krock et al., 2009), or the defects seen are less severe than those seen in anterograde IFT gene mutants (Tsujikawa and Malicki, 2004). There is evidence that some IFT genes can function outside of cilia (Cong et al., 2014; Delaval et al., 2011; Finetti et al., 2009; Finetti et al., 2014), so some of the differences seen in phenotypes of different IFT gene mutants may be due to roles of these genes outside of cilia rather than their function in generating and maintaining cilia.

In contrast to what was previously seen in photoreceptors, we found evidence of hair cells undergoing apoptosis in comparable numbers in the lateral line of both *ift88* and *dync2h1* gene mutants (Figures 1 & 2). This suggests the requirement of *ift88* for hair cell survival is due to its role more globally in cilia formation and maintenance rather than something gene-specific. It also shows that IFT genes are important for the survival of both inner ear and lateral line hair cells of zebrafish. However, the number of apoptotic cells we saw did not fully account for the hair cell number differences seen in these mutants. We usually saw between 3-4 fewer hair cells/neuromast in IFT gene mutants at 5 and 6 dpf, whereas we would only see 1-2 hair cells undergoing apoptosis in a small subset of lateral line neuromasts from 4-6 dpf (Figures 1 & 2). There could be a few reasons for this discrepancy. As we are only fixing at distinct time points we are certainly not seeing all the hair cells that will undergo apoptosis. Additionally, it is possible that some hair cells are dying by mechanisms other than apoptosis, such as necrosis, and thus are not staining positive for cleaved caspase-3. It is also possible that cilia are impacting hair cell numbers by affecting something other than hair cell survival. Though as discussed below we did not see obvious impacts on hair cell development.

We also found evidence that disruption in mitochondria may play a role in the hair cell death seen in IFT gene mutants. This is consistent with data in other cell types showing disrupted mitochondria activity and/or morphology in cilia gene mutants (Fujii et al., 2021; Lee et al., 2018; Lee et al., 2021) and that these mitochondria disruptions are linked to cell death (Bae et al., 2019; Lee et al., 2021). We saw a small but significant decrease in mitochondrial membrane potential in both IFT gene mutants (Figure 3). We also saw that treatment with RU360, a mitochondrial uniporter inhibitor, led to a decrease in hair cell number in wild-type siblings but not IFT gene mutants (Figure 4). The mitochondrial uniporter functions to allow the mitochondria to uptake calcium from the cytoplasm (Bernardi, 1999; Gunter and Gunter, 1994; Kirichok et al., 2004), and plays key roles in cell death (Demaurex and Distelhorst, 2003; Scorrano et al., 2003). Our results are consistent with a model where mitochondrial uniporter function is impaired in IFT gene mutants and this leads to cell death, though they do not prove this model definitively. Future work is warranted to more thoroughly investigate how mitochondria are impacted in the hair cells of cilia gene mutants and what role this plays in hair cell death.

It is worth noting that decreases in hair cell activity can lead to reduced mitochondrial membrane (Pickett et al., 2018) and changes in mitochondria architecture (McQuate et al., 2023). Hair cells of IFT gene mutants show reduced uptake of the dye FM1-43 (Stawicki et al., 2016; Stawicki et al., 2019). As this rapid FM1-43 uptake is known to be dependent on hair cell activity (Gale et al., 2001; Meyers et al., 2003; Seiler and Nicolson, 1999) this suggests hair cell activity is reduced in these mutants.

Experiments looking at calcium imaging in response to water jet stimulation of lateral line hair cells also suggest hair cell activity is reduced in *ift88* gene mutants (Kindt et al., 2012). The reductions in mitochondrial membrane potential we observe in IFT gene mutants are not as dramatic as those observed in *cdh23* mutants (Pickett et al., 2018), which may be because *cdh23* mutants show a complete loss of hair cell activity (Nicolson et al., 1998; Seiler and Nicolson, 1999) whereas hair cell activity is only reduced but not eliminated in IFT gene mutants (Kindt et al., 2012; Stawicki et al., 2016; Stawicki et al., 2019). Thus it is not clear if the mitochondria defects we see in IFT gene mutants are due to direct impacts of the cilia on the mitochondria or indirect impacts due to changes in hair cell activity.

Defects in proliferation during development are another way by which IFT gene mutants could impact hair cell number. This is commonly seen in neuronal development (Amador-Arjona et al., 2011; Breunig et al., 2008; Chizhikov et al., 2007; Han et al., 2008; Lepanto et al., 2016; Pruski et al., 2019; Spassky et al., 2008; Tong et al., 2014). Our results suggest that *dync2h1,* and thus cilia in general, do not play a role in hair cell proliferation during development, however, it is not clear if *ift88* is playing a role. To investigate whether decreased proliferation during hair cell development is responsible for the reduction in hair cell number seen in IFT gene mutants we treated fish with EdU from 3-4 dpf and then hair cells were labeled at 5dpf. These time points were chosen as decreases in hair cell number in *dync2h1* mutants are first observed at 5dpf (Stawicki et al., 2016)(Figure 2). When doing this we failed to see any significant differences in the number of EdU-positive hair cells in either the *ift88* or *dync2h1* mutants, though the *ift88* mutants did show slightly fewer EdU-positive hair cells/neuromast, with the average being 4.4 for wild-type siblings and 3.5 for mutants (Figure 5 A-C). However, during the regeneration experiments when fish were treated with EdU from 4-5 dpf and then hair cells labeled at 7 dpf, we did see a significant difference in the number of EdU-positive hair cells/neuromast in the EM control group for the *ift88* mutants which had not undergone regeneration, with the average being 4 for wild-type siblings and 2 for mutants (Figure 8). One possibility for this disparity is that the role of IFT genes in proliferation during hair cell development varies by age, however, unlike *dync2h1* mutants, *ift88* mutants are already showing significant decreases in hair cell number at 4 dpf (Figure 1), so if proliferation was playing a role in that difference one would expect it to be present early in development. Alternatively, as the regeneration experiment labeled hair cells 2 days rather than 1 day after EdU labeling, and we know there is hair cell death in these mutants, there may be more death of new hair cells in those experiments thus lowering the EdU-positive hair cell counts. Upon repeating the developmental EdU experiments and fixing 2 days after EdU labeling rather than 1 day (thus fish were labelled with EdU from 3-4 dpf and fixed at 6dpf) we did see a significant reduction in the number of EdU positive hair cells in *ift88* mutants (Figure 5D) suggesting this is what is happening.

Another way by which hair cell number could be changed in IFT gene mutants would be defects in hair cell differentiation. To investigate this we looked at supporting cell number reasoning that if hair cells were not differentiating we may see an increase in cells kept at a supporting cell fate. However, we failed to see any differences in supporting cell number between wild-type siblings and IFT gene mutants for either mutant (Figure 6). Thus, this was another situation where we failed to see impacts on neuromast or hair cell development in IFT gene mutants. In addition to suggesting hair cell differentiation may not be altered these results also further support the theory that there are not widespread proliferation defects during hair cell development in IFT gene mutants, as these would presumably decrease supporting cell number as well as hair cell number. However, from our data we cannot rule out defects in hair cell differentiation that do not change supporting cell number.

While we did not see consistent evidence for defects in proliferation during hair cell development across the two different IFT mutants, both mutants did show significant decreases in proliferation and subsequent hair cell number during regeneration (Figures 7 and 8). However, as it is known that neuromasts generally regenerate back to their starting size (Ma et al., 2008), it is not clear how much of this is due to impairment in the regenerative process versus the smaller starting size of neuromasts in IFT mutants.

Indeed, mutants that had undergone hair cell regeneration actually had more hair cells, and thus closer hair cell numbers to their wild-type siblings, than those that had not undergone regeneration (Figure 7). This would fit with a model where hair cells are dying over time creating the difference in hair cell number between wild-type siblings and mutants, as the regeneration response would have led to the birth of new hair cells that would not yet have had time to undergo apoptosis.

Overall our results suggest that the decrease in hair cell number in *dync2h1* mutants seen in normal situations is in large part due to hair cell death. We do not see evidence of defects in proliferation during development in these mutants. Additionally, significant hair cell number differences are not seen until 5 dpf in these mutants (Figure 2) (Stawicki et al., 2016) which would fit more with a model of hair cell death than impaired initial development. The more dramatic differences in hair cell number in fish that have not undergone regeneration than those that did also fit this model as discussed above. In contrast to this, *ift88* may be playing other roles in controlling hair cell number due to the earlier observed decrease in hair cell number (Figure 1) and potential decreases in proliferation in situations where regeneration is not occurring (Figures 4 and 8). It also appears IFT genes are not required for hair cell regeneration as it can still occur in mutants, but regeneration and proliferation of cells during regeneration is reduced in these mutants (Figures 7 and 8). This could be due to an impairment in the regenerative process itself or reductions in initial hair cell number.

## ACKNOWLEDGEMENTS

This work was supported by two Lafayette College Academic Research Committee faculty research grants to TMS, a Lafayette College Academic Research Committee student research grant to JG, and the Lafayette College Neuroscience program. We thank Erin Jiminez for technical advice and Amy Badillo for assistance with zebrafish care.

## COMPETING INTERESTS

The authors have no competing interests to declare

## Notes

### Competing Interest Statement

The authors have declared no competing interest.

### Summary of Updates

Additional data has been added looking at support cell number and number of EdU positive hair cells during development using different timing paradigms.

## REFERENCES

1. Abrams, S. R. and Reiter, J. F. (2021). Ciliary Hedgehog signaling regulates cell survival to build the facial midline. eLife 10, e68558.

2. Alexandre, D. and Ghysen, A. (1999). Somatotopy of the lateral line projection in larval zebrafish. Proc. Natl. Acad. Sci. 96, 7558–7562.

3. Amador-Arjona, A., Elliott, J., Miller, A., Ginbey, A., Pazour, G. J., Enikolopov, G., Roberts, A. J. and Terskikh, A. V. (2011). Primary cilia regulate proliferation of amplifying progenitors in adult hippocampus: implications for learning and memory. J. Neurosci. Off. J. Soc. Neurosci. 31, 9933–9944.

4. Bae, J.-E., Kang, G. M., Min, S. H., Jo, D. S., Jung, Y.-K., Kim, K., Kim, M.-S. and Cho, D.-H. (2019). Primary cilia mediate mitochondrial stress responses to promote dopamine neuron survival in a Parkinson’s disease model. Cell Death Dis. 10, 952.

5. Bernardi, P. (1999). Mitochondrial Transport of Cations: Channels, Exchangers, and Permeability Transition. Physiol. Rev. 79, 1127–1155.

6. Blanco-Sánchez, B., Clément, A., Fierro Junior, J., Washbourne, P. and Westerfield, M. (2014). Complexes of Usher proteins preassemble at the endoplasmic reticulum and are required for trafficking and ER homeostasis. Dis. Model. Mech. 7, 547–559.

7. Boubakri, M., Chaya, T., Hirata, H., Kajimura, N., Kuwahara, R., Ueno, A., Malicki, J., Furukawa, T. and Omori, Y. (2016). Loss of iY122, a Retrograde IFT Complex Component, Leads to Slow, Progressive Photoreceptor Degeneration Due to Inefficient Opsin Transport. J. Biol. Chem. 291, 24465–24474.

8. Breunig, J. J., Sarkisian, M. R., Arellano, J. I., Morozov, Y. M., Ayoub, A. E., Sojitra, S., Wang, B., Flavell, R. A., Rakic, P. and Town, T. (2008). Primary cilia regulate hippocampal neurogenesis by mediating sonic hedgehog signaling. Proc. Natl. Acad. Sci. 105, 13127– 13132.

9. Chizhikov, V. V., Davenport, J., Zhang, Q., Shih, E. K., Cabello, O. A., Fuchs, J. L., Yoder, B. K. and Millen, K. J. (2007). Cilia Proteins Control Cerebellar Morphogenesis by Promoting Expansion of the Granule Progenitor Pool. J. Neurosci. 27, 9780–9789.

10. Cong, E. H., Bizet, A. A., Boyer, O., Woerner, S., Gribouval, O., Filhol, E., Arrondel, C., Thomas, S., Silbermann, F., Canaud, G., et al. (2014). A homozygous missense mutation in the ciliary gene TTC21B causes familial FSGS. J. Am. Soc. Nephrol. JASN 25, 2435–2443.

11. Cullen, C. L., O’Rourke, M., Beasley, S. J., Auderset, L., Zhen, Y., Pepper, R. E., Gasperini, R. and Young, K. M. (2021). Kif3a deletion prevents primary cilia assembly on oligodendrocyte progenitor cells, reduces oligodendrogenesis and impairs fine motor function. Glia 69, 1184–1203.

12. Daudet, N., Gibson, R., Shang, J., Bernard, A., Lewis, J. and Stone, J. (2009). Notch regulation of progenitor cell behavior in quiescent and regenerating auditory epithelium of mature birds. Dev. Biol. 326, 86–100.

13. Delaval, B., Bright, A., Lawson, N. D. and Doxsey, S. (2011). The cilia protein IFT88 is required for spindle orientation in mitosis. Nat. Cell Biol. 13, 461–468.

14. Demaurex, N. and Distelhorst, C. (2003). Apoptosis--the Calcium Connection. Science 300, 65– 67.

15. Doerre, G. and Malicki, J. (2002). Genetic analysis of photoreceptor cell development in the zebrafish retina. Mech. Dev. 110, 125–138.

16. Ezratty, E. J., Stokes, N., Chai, S., Shah, A. S., Williams, S. E. and Fuchs, E. (2011). A Role for the Primary Cilium in Notch Signaling and Epidermal Differentiation during Skin Development. Cell 145, 1129–1141.

17. Ferraro, S., Gomez-Montalvo, A. I., Olmos, R., Ramirez, M. and Lamas, M. (2015). Primary cilia in rat mature Müller glia: downregulation of IFT20 expression reduces sonic hedgehog-mediated proliferation and dedifferentiation potential of Müller glia primary cultures. Cell. Mol. Neurobiol. 35, 533–542.

18. Fined, F., Paccani, S. R., Riparbelli, M. G., Giacomello, E., Perined, G., Pazour, G. J., Rosenbaum, J. L. and Baldari, C. T. (2009). Intraflagellar transport is required for polarized recycling of the TCR/CD3 complex to the immune synapse. Nat. Cell Biol. 11, 1332–1339.

19. Fined, F., Patrussi, L., Masi, G., Onnis, A., Galgano, D., Lucherini, O. M., Pazour, G. J. and Baldari, C. T. (2014). Specific recycling receptors are targeted to the immune synapse by the intraflagellar transport system. J. Cell Sci. 127, 1924–37.

20. Fujii, R., Hasegawa, S., Maekawa, H., Inoue, T., Yoshioka, K., Uni, R., Ikeda, Y., Nangaku, M. and Inagi, R. (2021). Decreased IFT88 expression with primary cilia shortening causes mitochondrial dysfunction in cisplatin-induced tubular injury. Am. J. Physiol. Renal Physiol. 321, F278–F292.

21. Gale, J. E., Marcod, W., Kennedy, H. J., Kros, C. J. and Richardson, G. P. (2001). FM1-43 dye behaves as a permeant blocker of the hair-cell mechanotransducer channel. J. Neurosci. 21, 7013–7025.

22. Gorivodsky, M., Mukhopadhyay, M., Wilsch-Braeuninger, M., Phillips, M., Teufel, A., Kim, C., Malik, N., Huttner, W. and Westphal, H. (2009). Intraflagellar transport protein 172 is essential for primary cilia formation and plays a vital role in paierning the mammalian brain. Dev. Biol. 325, 24–32.

23. Graf, M., Chakchouk, I., Ma, Q., Bensaid, M., DeSmidt, A., Turki, N., Yan, D., Baanannou, A., Mittal, R., Driss, N., et al. (2015). A missense mutation in DCDC2 causes human recessive deafness DFNB66, likely by interfering with sensory hair cell and supporting cell cilia length regulation. Hum. Mol. Genet. 24, 2482–2491.

24. Grisanf, L., Revenkova, E., Gordon, R. E. and Iomini, C. (2016). Primary cilia maintain corneal epithelial homeostasis by regulation of the Notch signaling pathway. Dev. Camb. Engl. 143, 2160–2171.

25. Gross, J. M., Perkins, B. D., Amsterdam, A., Egaña, A., Darland, T., Matsui, J. I., Sciascia, S., Hopkins, N. and Dowling, J. E. (2005). Identification of Zebrafish Insertional Mutants With Defects in Visual System Development and Function. GeneGcs 170, 245–261.

26. Gunter, K. K. and Gunter, T. E. (1994). Transport of calcium by mitochondria. J. Bioenerg. Biomembr. 26, 471–485.

27. Han, Y.-G., Spassky, N., Romaguera-Ros, M., Garcia-Verdugo, J.-M., Aguilar, A., Schneider-Maunoury, S. and Alvarez-Buylla, A. (2008). Hedgehog signaling and primary cilia are required for the formation of adult neural stem cells. Nat. Neurosci. 11, 277–284.

28. Hernández, P. P., Olivari, F. A., Sarrazin, A. F., Sandoval, P. C. and Allende, M. L. (2007). Regeneration in zebrafish lateral line neuromasts: Expression of the neural progenitor cell marker sox2 and proliferation-dependent and-independent mechanisms of hair cell renewal. Dev. Neurobiol. 67, 637–654.

29. Hudak, L. M., Lunt, S., Chang, C.-H., Winkler, E., Flammer, H., Lindsey, M. and Perkins, B. D. (2010). The intraflagellar transport protein iY80 is essential for photoreceptor survival in a zebrafish model of jeune asphyxiating thoracic dystrophy. Invest. Ophthalmol. Vis. Sci. 51, 3792–3799.

30. Kindt, K. S., Finch, G. and Nicolson, T. (2012). Kinocilia mediate mechanosensitivity in developing zebrafish hair cells. Dev. Cell 23, 329–341.

31. Kirichok, Y., Krapivinsky, G. and Clapham, D. E. (2004). The mitochondrial calcium uniporter is a highly selective ion channel. Nature 427, 360–364.

32. Kitami, M., Yamaguchi, H., Ebina, M., Kaku, M., Chen, D. and Komatsu, Y. (2019). IFT20 is required for the maintenance of cartilaginous matrix in condylar cartilage. Biochem. Biophys. Res. Commun. 509, 222–226.

33. Krock, B. L., Mills-Henry, I. and Perkins, B. D. (2009). Retrograde intraflagellar transport by cytoplasmic dynein-2 is required for outer segment extension in vertebrate photoreceptors but not arrestin translocation. Invest. Ophthalmol. Vis. Sci. 50, 5463–71.

34. Lee, J., Yi, S., Won, M., Song, Y. S., Yi, H.-S., Park, Y. J., Park, K. C., Kim, J. T., Chang, J. Y., Lee, M. J., et al. (2018). Loss-of-function of IFT88 determines metabolic phenotypes in thyroid cancer. Oncogene 37, 4455–4474.

35. Lee, J., Park, K. C., Sul, H. J., Hong, H. J., Kim, K.-H., Kero, J. and Shong, M. (2021). Loss of primary cilia promotes mitochondria-dependent apoptosis in thyroid cancer. Sci. Rep. 11, 4181.

36. Lepanto, P., Davison, C., Casanova, G., Badano, J. L. and Zolessi, F. R. (2016). Characterization of primary cilia during the differentiation of retinal ganglion cells in the zebrafish. Neural Develop. 11, 10.

37. Li, X., Yang, S., Han, L., Mao, K. and Yang, S. (2020a). Ciliary IFT80 is essential for intervertebral disc development and maintenance. FASEB J. Off. Publ. Fed. Am. Soc. Exp. Biol. 34, 6741– 6756.

38. Li, X., Lu, Q., Peng, Y., Geng, F., Shao, X., Zhou, H., Cao, Y. and Zhang, R. (2020b). Primary cilia mediate Klf2-dependant Notch activation in regenerating heart. Protein Cell 11, 433– 445.

39. Lin-Jones, J., Parker, E., Wu, M., Knox, B. E. and Burnside, B. (2003). Disruption of kinesin II function using a dominant negative-acting transgene in Xenopus laevis rods results in photoreceptor degeneration. Invest. Ophthalmol. Vis. Sci. 44, 3614–3621.

40. Liu, Z., Tu, H., Kang, Y., Xue, Y., Ma, D., Zhao, C., Li, H., Wang, L. and Liu, F. (2019). Primary cilia regulate hematopoietic stem and progenitor cell specification through Notch signaling in zebrafish. Nat. Commun. 10, 1839.

41. Lopes, V. S., Jimeno, D., Khanobdee, K., Song, X., Chen, B., Nusinowitz, S. and Williams, D. S. (2010). Dysfunction of heterotrimeric kinesin-2 in rod photoreceptor cells and the role of opsin mislocalization in rapid cell death. Mol. Biol. Cell 21, 4076–4088.

42. Ma, E. Y., Rubel, E. W. and Raible, D. W. (2008). Notch Signaling Regulates the Extent of Hair Cell Regeneration in the Zebrafish Lateral Line. J. Neurosci. 28, 2261–2273.

43. McQuate, A., Knecht, S. and Raible, D. W. (2023). Activity regulates a cell type-specific mitochondrial phenotype in zebrafish lateral line hair cells. eLife 12, e80468.

44. Meyers, J. R., MacDonald, R. B., Duggan, A., Lenzi, D., Standaert, D. G., Corwin, J. T. and Corey, D. P. (2003). Lighting up the senses: FM1-43 loading of sensory cells through nonselective ion channels. J. Neurosci. 23, 4054–4065.

45. Mill, P., Christensen, S. T. and Pedersen, L. B. (2023). Primary cilia as dynamic and diverse signalling hubs in development and disease. Nat. Rev. Genet. 24, 421–441.

46. Mizutari, K., Fujioka, M., Hosoya, M., Bramhall, N., Okano, H. J., Okano, H. and Edge, A. S. B. (2013). Notch inhibition induces cochlear hair cell regeneration and recovery of hearing aYer acoustic trauma. Neuron 77, 58–69.

47. Nicolson, T., Rüsch, A., Friedrich, R. W., Granato, M., Ruppersberg, J. P. and Nüsslein-Volhard, C. (1998). Genetic Analysis of Vertebrate Sensory Hair Cell Mechanosensation: the Zebrafish Circler Mutants. Neuron 20, 271–283.

48. Noda, K., Kitami, M., Kitami, K., Kaku, M. and Komatsu, Y. (2016). Canonical and noncanonical intraflagellar transport regulates craniofacial skeletal development. Proc. Natl. Acad. Sci. U. S. A. 113, E2589–2597.

49. Palla, A. R., Hilgendorf, K. I., Yang, A. V., Kerr, J. P., Hinken, A. C., Demeter, J., Krak, P., Mooney, N. A., Yucel, N., Burns, D. M., et al. (2022). Primary cilia on muscle stem cells are critical to maintain regenerative capacity and are lost during aging. Nat. Commun. 13, 1439.

50. Pazour, G. J., Baker, S. A., Deane, J. A., Cole, D. G., Dickert, B. L., Rosenbaum, J. L., Witman, G. B. and Besharse, J. C. (2002). The intraflagellar transport protein, IFT88, is essential for vertebrate photoreceptor assembly and maintenance. J. Cell Biol. 157, 103–13.

51. Pickett, S. B., Thomas, E. D., Sebe, J. Y., Linbo, T., Esterberg, R., Hailey, D. W. and Raible, D. W. (2018). Cumulative mitochondrial activity correlates with ototoxin susceptibility in zebrafish mechanosensory hair cells. eLife 7, e38062.

52. Pruski, M., Hu, L., Yang, C., Wang, Y., Zhang, J.-B., Zhang, L., Huang, Y., Rajnicek, A. M., St Clair, D., McCaig, C. D., et al. (2019). Roles for IFT172 and Primary Cilia in Cell Migration, Cell Division, and Neocortex Development. Front. Cell Dev. Biol. 7, 287.

53. Raible, D. W. and Kruse, G. J. (2000). Organization of the lateral line system in embryonic zebrafish. J. Comp. Neurol. 421, 189–198.

54. Reers, M., Smith, T. W. and Chen, L. B. (1991). J-aggregate formation of a carbocyanine as a quantitative fluorescent indicator of membrane potential. Biochemistry 30, 4480–4486.

55. Reiter, J. F. and Leroux, M. R. (2017). Genes and molecular pathways underpinning ciliopathies. Nat. Rev. Mol. Cell Biol. 18, 533–547.

56. Ryan, S., Willer, J., Marjoram, L., Bagwell, J., Mankiewicz, J., Leshchiner, I., Goessling, W., Bagnat, M. and Katsanis, N. (2013). Rapid identification of kidney cyst mutations by whole exome sequencing in zebrafish. Development 140, 4445–51.

57. Santra, P. and Amack, J. D. (2021). Loss of vacuolar-type H+-ATPase induces caspase-independent necrosis-like death of hair cells in zebrafish neuromasts. Dis. Model. Mech. 14, dmm048997.

58. Scorrano, L., Oakes, S. A., Opferman, J. T., Cheng, E. H., Sorcinelli, M. D., Pozzan, T. and Korsmeyer, S. J. (2003). BAX and BAK Regulation of Endoplasmic Reticulum Ca2+: A Control Point for Apoptosis. Science 300, 135–139.

59. Seiler, C. and Nicolson, T. (1999). Defective calmodulin-dependent rapid apical endocytosis in zebrafish sensory hair cell mutants. J. Neurobiol. 41, 424–434.

60. Song, B., Haycrak, C. J., Seo, H., Yoder, B. K. and Serra, R. (2007). Development of the post-natal growth plate requires intraflagellar transport proteins. Dev. Biol. 305, 202–216.

61. Spassky, N., Han, Y.-G., Aguilar, A., Strehl, L., Besse, L., Laclef, C., Ros, M. R., Garcia-Verdugo, J. M. and Alvarez-Buylla, A. (2008). Primary cilia are required for cerebellar development and Shh-dependent expansion of progenitor pool. Dev. Biol. 317, 246–259.

62. Stawicki, T. M., Owens, K. N., Linbo, T., Reinhart, K. E., Rubel, E. W. and Raible, D. W. (2014). The zebrafish merovingian mutant reveals a role for pH regulation in hair cell toxicity and function. Dis. Model. Mech. 7, 847–856.

63. Stawicki, T. M., Hernandez, L., Esterberg, R., Linbo, T., Owens, K. N., Shah, A. N., Thapa, N., Roberts, B., Moens, C. B., Rubel, E. W., et al. (2016). Cilia-Associated Genes Play Differing Roles in Aminoglycoside-Induced Hair Cell Death in Zebrafish. G3 GenesGenomesGeneGcs 6, 2225–2235.

64. Stawicki, T. M., Linbo, T., Hernandez, L., Parkinson, L., Bellefeuille, D., Rubel, E. W. and Raible, D. W. (2019). The role of retrograde intraflagellar transport genes in aminoglycoside-induced hair cell death. Biol. Open 8, bio038745.

65. Tao, D., Zhang, L., Ding, Y., Tang, N., Xu, X., Li, G., Niu, P., Yue, R., Wang, X., Shen, Y., et al. (2023). Primary cilia support cartilage regeneration aYer injury. Int. J. Oral Sci. 15, 22.

66. Taschner, M. and Lorentzen, E. (2016). The Intraflagellar Transport Machinery. Cold Spring Harb. Perspect. Biol. 8, a028092.

67. Thomas, E. D. and Raible, D. W. (2019). Distinct progenitor populations mediate regeneration in the zebrafish lateral line. eLife 8,.

68. Thomas, E. D., Cruz, I. A., Hailey, D. W. and Raible, D. W. (2015). There and back again: development and regeneration of the zebrafish lateral line system. Wiley Interdiscip. Rev. Dev. Biol. 4, 1–16.

69. Tong, C. K., Han, Y.-G., Shah, J. K., Obernier, K., Guinto, C. D. and Alvarez-Buylla, A. (2014). Primary cilia are required in a unique subpopulation of neural progenitors. Proc. Natl. Acad. Sci. U. S. A. 111, 12438–12443.

70. Tsujikawa, M. and Malicki, J. (2004). Intraflagellar Transport Genes Are Essential for Differentiation and Survival of Vertebrate Sensory Neurons. Neuron 42, 703–716.

71. Wang, B., Zhang, Y., Dong, H., Gong, S., Wei, B., Luo, M., Wang, H., Wu, X., Liu, W., Xu, X., et al. (2018). Loss of Tctn3 causes neuronal apoptosis and neural tube defects in mice. Cell Death Dis. 9, 520.

72. Warchol, M. E., Stone, J., Barton, M., Ku, J., Veile, R., Daudet, N. and Lovett, M. (2017). ADAM10 and γ-secretase regulate sensory regeneration in the avian vestibular organs. Dev. Biol. 428, 39–51.

73. Yuan, X., Liu, M., Cao, X. and Yang, S. (2019). Ciliary IFT80 regulates dental pulp stem cells differentiation by FGF/FGFR1 and Hh/BMP2 signaling. Int. J. Biol. Sci. 15, 2087–2099.

